# Chronotype asymmetry arises from stochastic sleep homeostasis under circadian entrainment

**DOI:** 10.64898/2026.02.18.706715

**Authors:** Nguyen Trong Nguyen, Vuong Hung Truong, Jihwan Myung

**Author notes:** Correspondence: Jihwan Myung, PhD.

## Abstract

**Study Objectives:** Chronotype, typically defined by the mid-sleep phase on free days, is often interpreted as a direct proxy for circadian phase. However, a fundamental paradox challenges this assumption: worldwide chronotype distributions show a heavy right skew, whereas circadian period distributions show a slight left skew. We investigated whether fast, noisy cortical processes in homeostatic sleep pressure dynamics could resolve this paradox.

**Methods:** Drawing upon the classical two-process model of sleep regulation, we simulated how interindividual differences in sleep homeostasis (Process S), combined with realistic circadian period distributions (Process C), interact to shape chronotype variability. We introduce stochasticity into sleep pressure build-up to test whether homeostatic variance generates the empirically observed right-skew in chronotype distributions.

**Results:** Exploring the parameter space reveals that stable sleep-wake cycles are constrained within physiological limits of entrainment in an Arnold tongue-like structure. While our simulations confirm a general positive correlation between intrinsic period and chronotype, the heavy right tail of the chronotype distribution arises from interindividual variability in homeostatic sleep dynamics operating near the unstable boundaries of entrainment. This mechanism preserves the individual-level correlation but inverts the population-level skew from left to right.

**Conclusions:** Behavioral chronotype emerges from the nonlinear interaction between fluctuating homeostatic sleep pressure and a hierarchically entrained circadian system. We caution against inferring the intrinsic circadian period solely from chronotype, particularly in clinical contexts where accurate assessment of circadian rhythmicity is critical for targeted therapies.

## 1. Introduction

Chronotypes reflect interindividual differences in daily temporal patterns of activity, typically expressed as preferences in sleep-wake timing (Roenneberg et al., 2003), with growing evidence linking chronotype to psychiatric vulnerability (Zou et al., 2022). Under circadian entrainment, these behavioral preferences can be interpreted as indicators of the phases of the endogenous circadian clock, which is determined by the intrinsic circadian period and its interaction with external time cues (Bordyugov et al., 2015). However, chronotype is defined behaviorally rather than physiologically, and therefore does not provide a direct readout of the circadian clock itself. Instead, it is a multidimensional construct shaped by multiple underlying cyclic processes, including sleep timing, eating behavior, body temperature, hormone secretion, and cognitive functions (Chauhan et al., 2023). Conceptually, chronotype can be operationalized as a phase of entrainment, for example, as the point of mid-sleep or midday, which depends on the molecular clock and downstream behavioral and physiological processes, and varies across different photoperiod conditions (Schmal et al., 2020).

The influence of the circadian clock on entrainment phase has long been expected to yield a linear relationship between circadian period length and chronotype. Evidence from a small sample (*n* = 17) study by Duffy et al. (2001) showed a strong correlation (*r* = 0.6), with morning types exhibiting shorter periods (24.1 h) than evening types (24.3 h). A more recent study with a larger sample size (*n* = 157) reported a similar, significant correlation (*r* = 0.41) (Crowley & Eastman, 2018; **Figure 1A**). Likewise, Brown et al. (2008) reported that morning types had slightly shorter circadian periods (24.33 ± 0.41 h) than late chronotypes (24.74 ± 0.32 h), with a moderate correlation between circadian period and entrained phase (Spearman ρ = 0.36). Additionally, Hasan et al. (2012) demonstrated consistently across *Per3* polymorphisms that morning types had a shorter *in vivo* circadian period than evening types, with a significant correlation, though not for *in vitro* periods.

**Figure 1.**
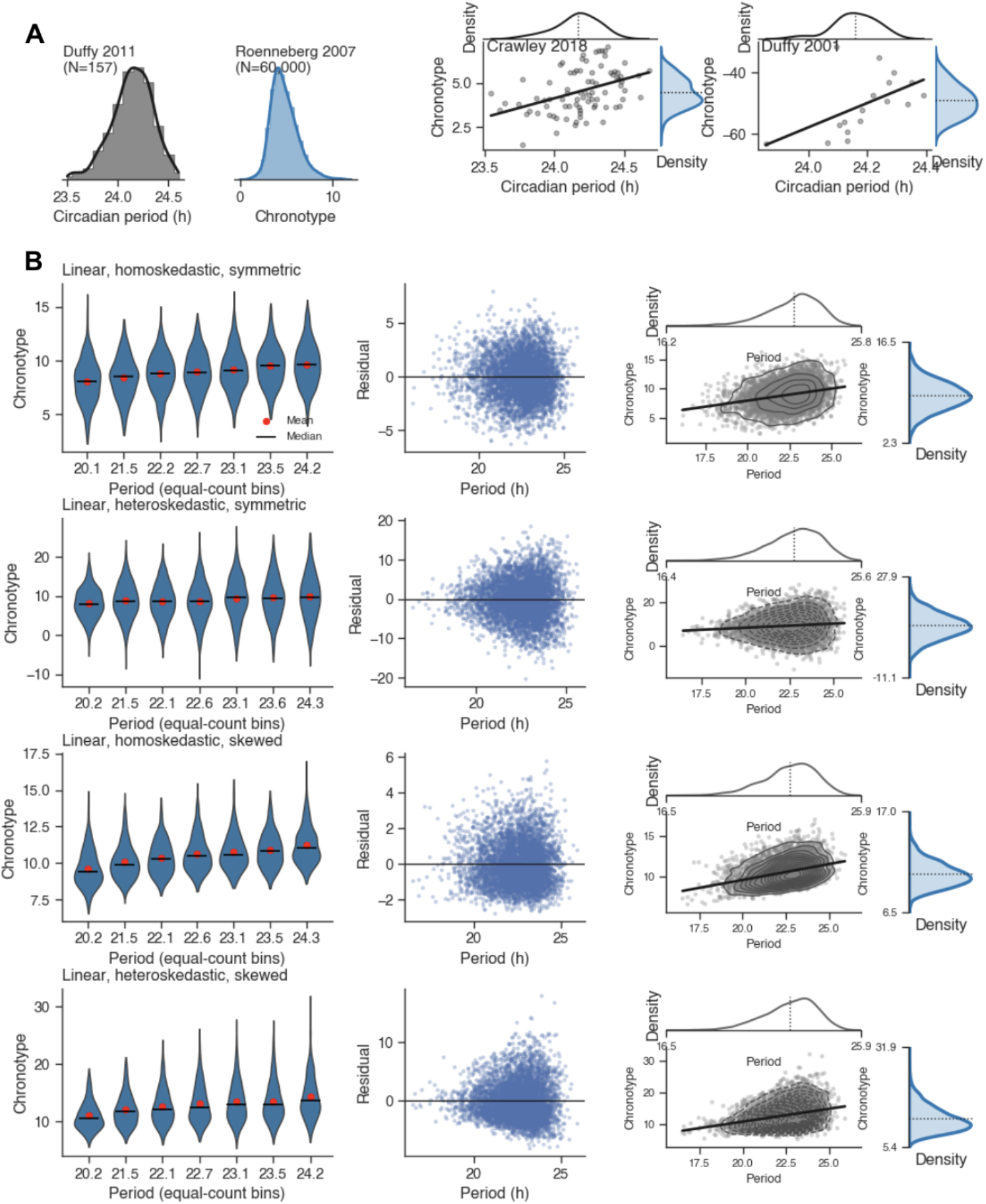
Empirical and simulated relationships between circadian period and chronotype. **(A)** Empirical distributions and associations between intrinsic circadian period and chronotype across large human datasets. Left: population distributions of intrinsic circadian period and chronotype in MSF, reconstructed from published data using smoothed density estimates (bars indicate original binned counts; lines indicate smoothed densities). Right: joint distributions illustrating the relationship between circadian period and chronotype for individual-level datasets (in MSF, Crowley & Eastman, 2018; in inverted MEQ, Duffy et al., 2011). Points represent individuals, solid lines indicate least-squares linear fits, and marginal densities are shown for both circadian period and chronotype. Notably, the circadian period exhibits a left-skewed distribution, whereas chronotype shows a right-skewed distribution, suggesting a deviation from strict linearity in their relationship. **(B)** Simulated examples illustrating how distributional properties influence observed period-chronotype relationships. Four rows correspond to linear relationships with increasing complexity: homoskedastic symmetric noise, heteroskedastic noise, skewed noise, and combined heteroskedastic and skewed noise. Columns show (left) binned chronotype distributions across equal-count period bins (violins), with bin means (red points) and medians (black bars); (middle) residuals from linear regression as a function of period; and (right) joint distributions highlighting the resulting shape of the period-chronotype association. These simulations demonstrate that apparent nonlinearities similar to those observed in (A) can arise from skewed, and particularly right-skewed noise, even when the underlying relationship is linear.

If this linear relationship were sufficient to explain individual differences, the population distribution of chronotype would be expected to mirror that of intrinsic circadian periods. However, while chronotype distributions are consistently right-skewed (Fischer et al., 2017; Roenneberg et al., 2007; Sládek et al., 2020), free-running circadian period distributions have been reported as left-skewed (Crowley & Eastman, 2018; Duffy et al., 2001, 2011), approximately Gaussian (Hiddinga et al., 1997; Wright et al., 2005), or right-skewed in small sample studies (Burgess & Eastman, 2008; Czeisler et al., 1999). This mismatch suggests that additional factors modulate the translation from intrinsic period to behavioral chronotype.

Sleep-wake timing reflects not only circadian phase, but also the homeostatic sleep drive as well as environmental cues such as light and social timing. Consequently, chronotype is typically assessed via behavioral measures of sleep-wake timing, such as self-reported schedules, questionnaires, sleep diaries, or actigraphy, rather than by direct physiological readouts of the clock. These measures primarily capture mid-sleep on free days (MSF), a robust proxy for circadian phase, as evidenced by its correlation with dim-light melatonin onset (DLMO), a gold-standard marker of circadian timing (Roenneberg et al., 2003). In the present study, we adopt this definition to assess chronotypes in simulated free-running conditions (i.e., in the absence of a light-dark cycle). By combining realistic distributions of intrinsic circadian periods with interindividual variability in sleep homeostasis, we aim to investigate the origins of the observed discrepancy between period and chronotype.

Our approach is grounded in the two-process model, which conceptualizes the sleep-wake cycle as two partially independent processes coupled to the circadian clock: the circadian rhythm (Process C) and the homeostatic sleep drive (Process S) (Borbély, 1982). Process C, expressed through physiological oscillations of body temperature and melatonin secretion, is regulated by the hypothalamic suprachiasmatic nucleus (SCN), whereas Process S reflects the accumulation of sleep pressure based on prior wakefulness, with slow-wave activity during sleep being primarily determined by time awake (Dijk & Czeisler, 1995). Under free-running conditions, these two processes can become partially uncoupled and run on their own periods, a phenomenon known as internal desynchronization.

Empirical evidence demonstrates that the circadian clock and the sleep-wake cycle can operate partially independently. Under free-running or isolated conditions, internal desynchronization can occur, with the circadian clock maintaining a relatively stable period while sleep-wake timing exhibits greater variability (Aschoff, 1965; Czeisler et al., 1980; Hashimoto et al., 1997; Yamanaka & Waterhouse, 2016). This partial independence underlines the need to consider both circadian and homeostatic influences when modeling chronotype.

To this end, we adopt the two-process model due to its robust empirical support, its ability to parameterize Process C while accounting for individual variability, and its capability to integrate circadian phase with homeostatic drive (Borbély et al., 2016). We extend the original framework by explicitly incorporating light as an external Zeitgeber, and create a hierarchy of three interacting oscillations: the sleep-wake process, the circadian clock, and the environmental light-dark cycle. This allows us to simulate how interindividual differences in sleep homeostasis and the circadian clock interact to shape chronotype variability under light entrainment (**Figure 2**). To account for the empirical right skew in chronotype distributions, we focus on potential asymmetries in homeostatic sleep processes. Variability in the rates of sleep pressure build-up and dissipation may differ across individuals, potentially introducing asymmetric timing of sleep onset and offset. In the present study, we investigate how such interindividual differences in homeostatic dynamics shape chronotype distributions.

**Figure 2.**
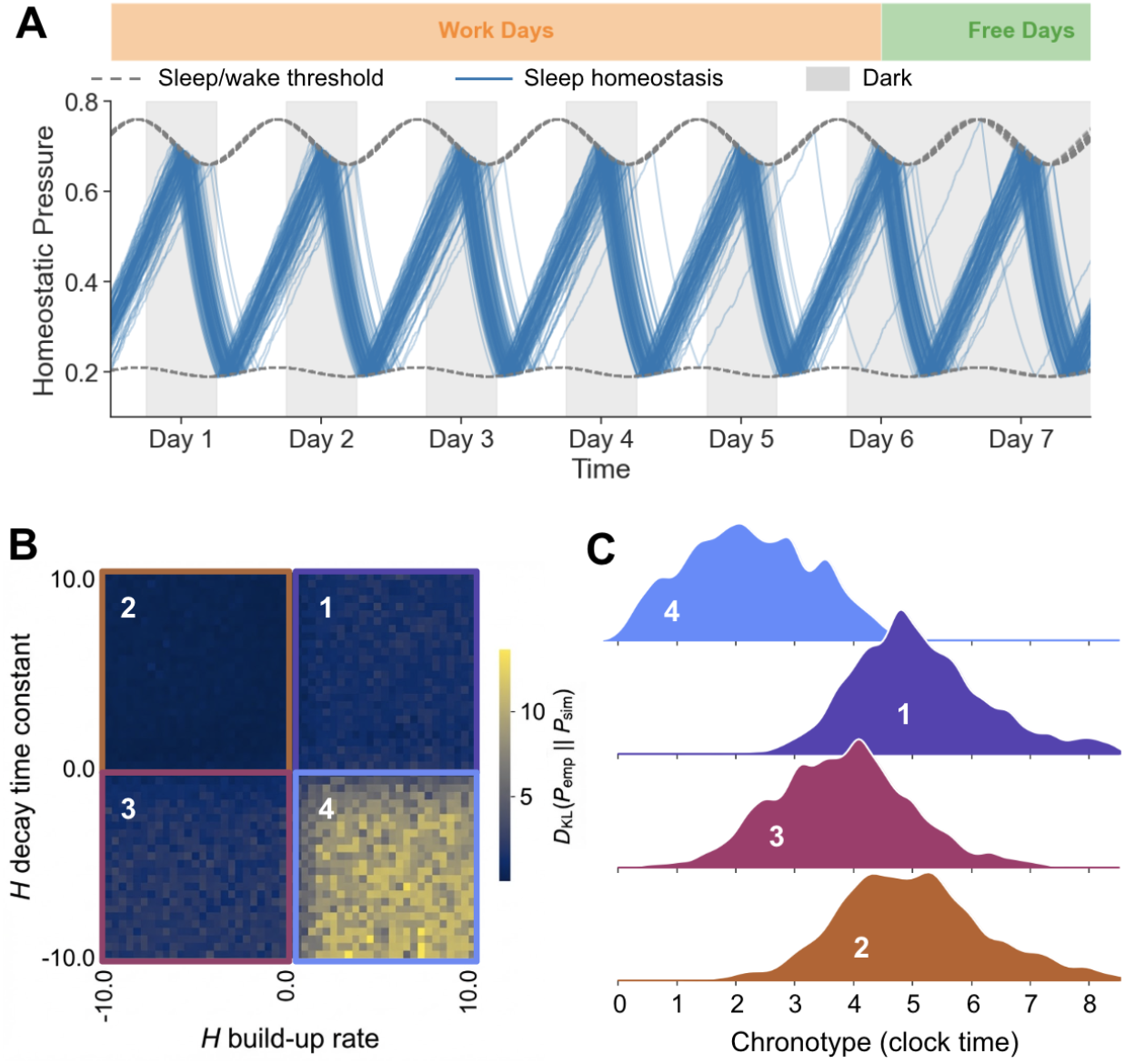
A linear-skewed generative model for chronotype. **(A)** Schematic of the modified two-process model incorporating stochastic homeostatic dynamics and external light forcing. The blue line represents the homeostatic sleep pressure trajectory. Sinusoidal boundaries define the sleep-wake thresholds, with the upper threshold triggering sleep onset and the lower threshold triggering awakening. Gray shaded regions indicate the release from external light forcing. The simulation proceeds for 14 days, with the final 7 days visualized to exclude initial transient drifts. **(B)** KL divergence landscape mapped against the distributional skewness of the sleep pressure build-up rate and the decay time constant. Lighter colors indicate lower information loss between simulated and empirical data, representing a better fit. Colored squares delineate the parameter space into four distinct quadrants. The global minimum for error is identified in the configuration characterized by rapid homeostatic accumulation and rapid dissipation. **(C)** Representative chronotype (as clock time) distributions resulting from the four homeostatic configurations in (B). Histogram colors correspond to the respective quadrants in the KL landscape. The chronotype derived from the second quadrant (red) exhibits the characteristic right skew, mean, and median observed in empirical data.

## 2. Methods

### Empirical data extraction

We extracted intrinsic circadian period and chronotype at both population and individual levels from published figures (Duffy et al., 2001, 2011; Crowley & Eastman, 2018; Fischer et al., 2017; Roenneberg et al., 2007; Sládek et al., 2020) using WebPlotDigitizer v5 (https://automeris.io/wpd/?v=5_2). Intrinsic circadian period (τ, h) was measured under constant or near-constant laboratory conditions. Chronotype was assessed either via morningness-eveningness questionnaires (MEQ) or by mid-sleep timing on free days (MSF). To enable comparison across datasets, MEQ scores were inverted so that higher values consistently indicated later chronotype (Zavada et al., 2005); no nonlinear transformations were applied.

### Generative linear model

To examine how distributional properties of noise influence the observed period-chronotype relationship, we simulated data under the model:

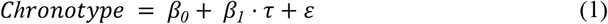

where *β*_*i*_ are the linear fit parameters, *τ* is the circadian period, and *ε* is stochastic noise. Four noise regimes were tested: (i) Homoskedastic, symmetric noise (Gaussian with constant variance); (ii) Heteroskedastic noise (variance increasing with circadian period); (iii) Skewed noise (right-skewed error distribution); (iv) Combined heteroskedastic and skewed noise. For each regime, chronotype distributions were examined across equal-count bins of circadian period.

### Two-process model

Following Daan et al. (1984), we modeled the upper (*U*) and lower (*L*) sleep thresholds as simple sinusoidal functions of the circadian phase *ϕ*, the amplitude *A*, and the mesors *M*_*U*_ and *M*_*L*_. We adopted an asymmetric threshold where the lower threshold amplitude is dampened to 20% of the upper threshold (i.e., *A*/5), and included a phase offset *α* applied after entrainment. This adjustment allowed us to shift the timing of the sleep thresholds relative to the stabilized circadian rhythm without altering the underlying entrainment dynamics:

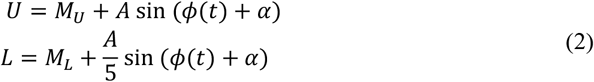

The phase *ϕ* is the solution of equation (4).

### Process C: circadian light entrainment

To analyze responses of clock dynamics to external perturbations, we abstracted away molecular details and modeled the circadian clock as a limit cycle oscillator, which, in the absence of external forcing, is governed solely by the intrinsic angular velocity *ω*:

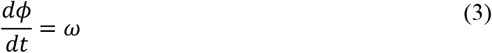

where *ω* = *2π/τ*, and *τ* is the circadian free-running period sampled from the empirical distribution of Duffy et al. (2011) (24.09 ± 0.2 h, mean ± standard deviation) via bootstrap-based kernel density estimation (1000 iterations). The median bootstrap density was used to construct a cumulative distribution function, from which estimated periods were drawn via inverse transform sampling.

For a continuously varying Zeitgeber, such as the daily variation of light intensity, we represented it as a sinusoidal wave with phase *ϕ*_*light*_ = (2*π*/24)*t* + *ϕ*_*lag*_, where *ϕ*_*lag*_ is a phase lag that adjusts to the local light-on time. To model entrainment, the influence of light on the internal clock was modeled by adding a phase-dependent interaction term gated by a sigmoid function that mimics phase-dependent sensitivity to light, *g*(*t*) = 1/(1 +exp (−10 ⋅ (sin *ϕ*_*light*_ − *θ*))), where *θ* = 0 during entrainment and *θ* = 20 during free-running, and *K*_*light*_ is the linear coupling constant (Myung et al., 2015):

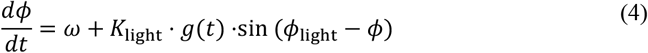

The resulting instantaneous phase *ϕ*(*t*) was substituted back into the threshold equations (2). Equation (4) was integrated using an explicit fourth-order Runge-Kutta (RK4) method with Numba just-in-time compilation at a step size of Δ*t* = 0.1 h.

### Simulation protocol

To obtain the chronotype, the model mimicked the real condition of social jet lag (Wittmann et al., 2006). After the initial stabilization period (7 days), it simulated “work day” light entrainment of a 24-h light-dark cycle for 5 days, followed by 2 days of “free day” free-running for assessment. To determine the individual’s final chronotype, we calculated the arithmetic mean of the mid-sleep times specifically during the free day period, capturing the interaction between biological preferences and social constraints (**Figure 2A**). The mid-sleep phase in clock time was estimated as chronotype, (*t*_*sleep*,_ mod 24 + *D*_*sleep*,_/2) mod 24, where *t*_*sleep*,_ is the sleep onset time and *D*_*sleep*,_ is the sleep duration.

### Process S: homeostatic sleep pressure with random walk

We modeled the homeostatic sleep pressure *H*(*t*) as a state variable that increases during wakefulness as a Brownian motion with drift and decays exponentially during sleep. The rate of sleep pressure accumulation during wake (*H*_*W*_) depends on neural activity, which varies with cognitive load and other faster timescale processes, and may therefore appear stochastic:

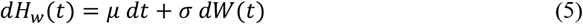

where *μ* is the mean drift rate, *σ* the noise intensity, and *dW*(*t*) the increment of a Wiener process, such that *dW* ∼ 𝒩(0, *dt*). The decay of sleep pressure (*H*_*s*_) during sleep is:

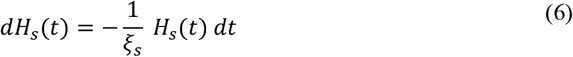

where *ξ*_*s*_ indicates the decay time constant of sleep pressure. For the numerical simulation, equations (5) and (6) were implemented using the discrete integration step:

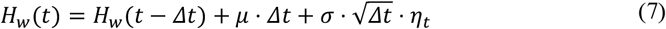

where *η*_*t*_∼ 𝒩(0, 1) i.i.d. and

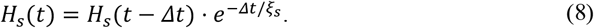

### Search for skewed distribution conditions

We quantified distributional similarity between simulated and empirical chronotype distributions by the Kullback-Leibler (KL) divergence. The empirical distributions were derived from large-scale surveys in the United States and Europe (mean: 4:36 AM, or 4.6 h; skew: 0.6; **Figure 1A**) (Fischer et al., 2017; Roenneberg et al., 2007). For each parameter configuration, we generated a simulated chronotype distribution from *n* = 400 individuals. Both the simulated and empirical distributions were converted to probability density functions using Gaussian kernel density estimation. The KL divergence was then calculated as:

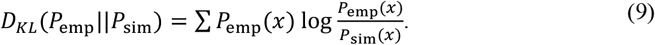

where *P*_emp_(*x*) is the empirical density and *P*_sim_(*x*) is the simulated density. To avoid numerical instability from zero bins, we added a small constant (10^−10^) before normalization. We used the KL divergence as an objective function for **Figures 2B** and **4B**.

Although KL divergence provides a robust measure of global distributional similarity and serves as our primary criterion for realism, it does not explicitly isolate dynamical stability boundaries. To better visualize the entrainment structure across parameter space, we defined a complementary composite error function (*Err*) that constrains key physiological summary statistics while allowing moderate deviations in higher-order shape such as variance, kurtosis, modality (e.g., unimodal versus multimodal structure). It quantifies the deviation in three key criteria: the median chronotype (*M*_sim_), the distribution skewness (*S*_sim_), and the average sleep duration (*D*_sim_):

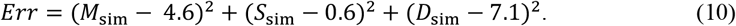

where the subscript sim denotes the simulated output. We targeted average sleep duration of 7.1 ± 0.4 h, a value consistently reported across diverse ethnic and age groups (Kocevska et al., 2020). Skewness was calculated as the third standardized moment to quantify the direction and magnitude of the tail:

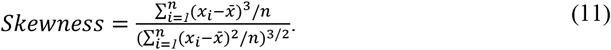

### Parameter search for chronotype distribution

We implemented a stepwise parameter search strategy to systematically isolate the determinants of chronotype. First, we analyzed the sensitivity of sleep homeostasis by fixing the circadian configuration to values established by Daan et al. (1984) that robustly reproduce monophasic sleep: circadian amplitude *A* = 0.05, mean upper threshold *M*_*U*_ = 0.71, and lower threshold *M*_*L*_ = 0.2. To capture biological heterogeneity, we sampled the homeostatic *μ* and *ξ*_*s*_ from theoretical skewed distributions, reflecting the non-Gaussian variability typical of physiological rates. To reduce complexity, we varied the skewness of the distribution while fixing the statistical moments: *ξ*_*s*_ centered at 5.5 (scale 0.5) and *μ* centered at 0.03 (scale 0.001), with noise intensity held constant at 0.005. Subsequently, we examined the circadian contribution by fixing the optimal homeostatic distribution and varying *A* and *M*_*U*_, while maintaining a constant *M*_*L*_ = 0.2. We quantified the dynamic impact of these configurations by mapping the threshold settings to the intrinsic sleep-wake cycle period, defined as the average interval between sleep events under static threshold conditions (*A* = 0). All base parameters are listed in **Table 1**.

**Table 1.**
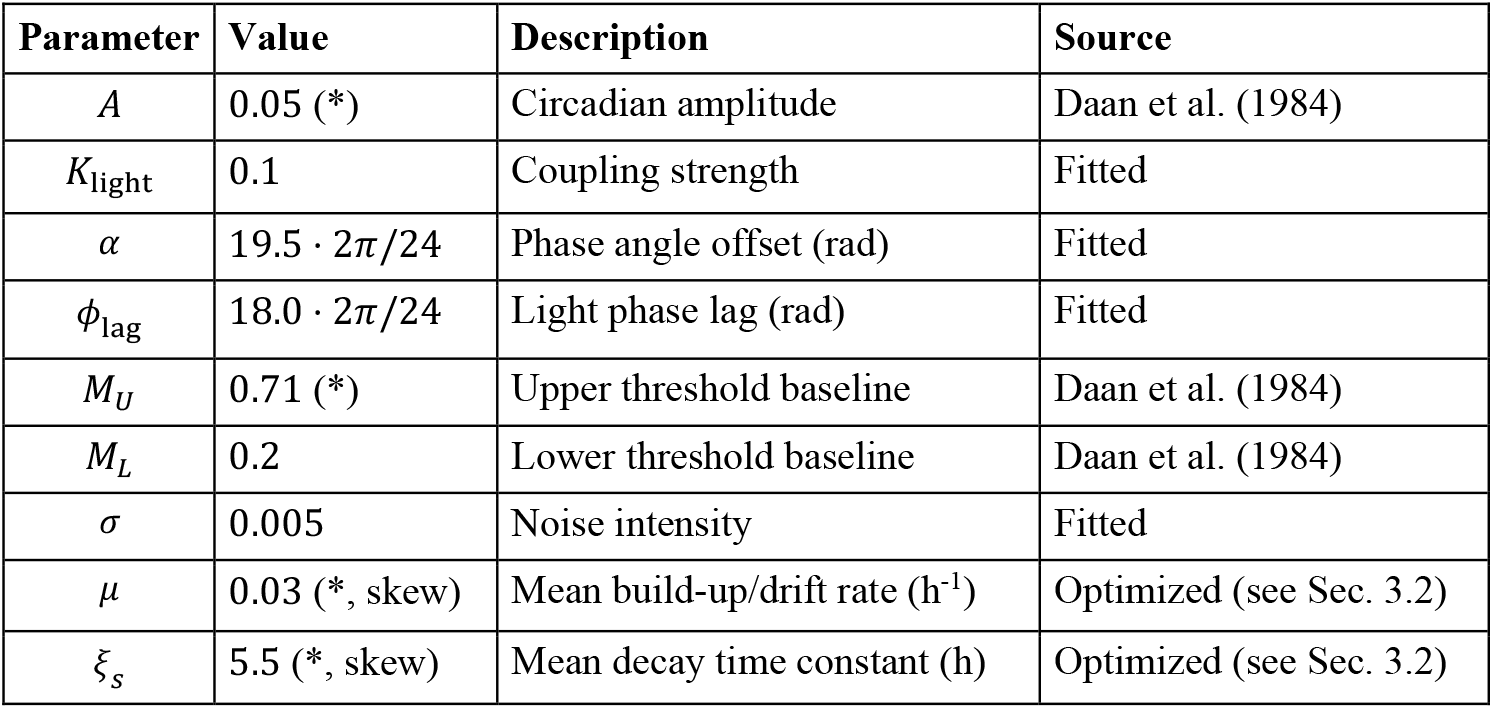
Base parameters of the model. Asterisk (*) indicates parameters that varied during parameter search.

## 3. Results

### 3.1. Circadian period and chronotype distributions show opposite skewness

Across empirical datasets, intrinsic circadian period exhibited a consistently left-skewed distribution, with a longer tail toward shorter periods. In contrast, chronotype showed a right-skewed distribution, characterized by a heavier tail toward later chronotypes. These opposing skewness patterns were robust across independent studies and measurement modalities. Despite a significant linear association between circadian period and chronotype at the individual level, joint distributions revealed systematic deviations from a simple linear trend. In particular, later chronotypes were disproportionately variable at longer circadian periods, producing an apparent curvature in the period-chronotype relationship (**Figure 1A**).

Simulated data with homoskedastic symmetric noise produced linear joint distributions and residuals consistent with classical regression assumptions (**Figure 1B**). Introducing heteroskedasticity alone increased variance at longer periods, but did not reproduce the asymmetric structure observed in empirical data. In contrast, right-skewed noise generated bin-wise divergences between means and medians, asymmetric residuals, and curved joint distributions closely resembling the empirical patterns. When skewness and heteroskedasticity were combined, these effects were amplified, while the direction and form of the skewness remained unchanged. Importantly, in all simulations the underlying generative relationship between circadian period and chronotype remained strictly linear. Thus, the observed nonlinear features emerged solely from the skewed distribution of noise.

### 3.2. Homeostatic parameter skewness shapes the chronotype distribution

We simulated sleep-wake dynamics under a social jet-lag protocol through distinct regimes of circadian entrainment under work day and free-running during free day (**Figure 2A**). Under work day conditions, the system maintained a stable phase relationship with the circadian process oscillating at 24 h period. Upon the transition to free days, phases drifted systematically as the circadian oscillator relaxed toward its natural frequency. We quantified intrinsic chronotype as the average mid-sleep phase position during the final two free days. This distinction clarifies the difference between conceptualizing chronotype as an internal phase relationship (*ψ*_int_) governed by the interaction of sleep homeostasis and the circadian oscillation, and as an external phase angle relative to light (*ψ*_ext_). While the circadian pacemaker remained stable across days, sleep-wake threshold crossings exhibited stochastic fluctuations, generating within-period variability in chronotype.

To interpret this variability mechanistically, we decomposed homeostatic dynamics into a drift component (*μ*) and a stochastic component (*σ*). The drift rate *μ* determines the mean time to sleep onset: a greater drift means Process S accumulates faster, reaching the upper threshold earlier and resulting in earlier sleep onset. The noise intensity *σ* does not change the mean onset time, but controls the spread around it. Critically, when sleep pressure build-up is formulated as a random walk, the duration of wakefulness becomes equivalent to a first-passage time, i.e., the time required for the trajectory *H*_*w*_(*t*) to reach the upper sleep threshold. If the threshold is static, first-passage times follow an inverse Gaussian (Wald) distribution, which is characterized by a distinctive positive skew. This is analogous to the decision time distribution in the drift-diffusion model (DDM) (Ratcliff & McKoon, 2008). Likewise, in Process S, symmetric fluctuations can generate asymmetric variability in sleep timing.

To systematically examine the origins of population asymmetry, we varied the skewness of the homeostatic build-up rate and the decay time constant while strictly constraining their mean and variance. Model fit was quantified using KL divergence to measure the information loss between the simulated and empirical chronotype distributions. The KL divergence heatmap (**Figure 2B**) revealed that the parameter space divides into distinct error landscapes centered around a Gaussian baseline of zero skew. The global minimum for error, representing the optimal biological fit, appears in the quadrant defined by a left-skewed build-up rate and a right-skewed decay constant. This configuration corresponds to bias toward rapid homeostatic accumulation and rapid dissipation. Notably, the gradient of the error landscape indicates that the shape of the population chronotype is significantly more sensitive to asymmetries in the decay constant than in the build-up rate. This highlights the dominant role of dissipation dynamics in shaping the stochastic boundaries of sleep episodes, as detailed in **Figure 2C**.

Traditionally, chronotype is modeled as a linear function of *τ*, a view grounded in deterministic oscillator theory (Pittendrigh, 1981). However, our simulations compel a reappraisal of this framework. While the fundamental positive correlation remains intact (*r* = 0.11, *p* = 0.01; **Figure 3A**), the linearity alone fails to account for the distributional inversion. **Figure 3B** demonstrates that the emergence of a right-skewed chronotype from a left-skewed period is driven by the asymmetric ‘noise’ of homeostatic sleep pressure. This interaction creates a state-dependent expansion of variance that is similar to the skewed heteroscedastic variance regime modeled in **Figure 1B. Figure 3C** illustrates the structured error profile, with variance peaking at intermediate periods and lower at the extremes. This confirms that chronotype is not a simple linear derivative of the clock, but is heavily modulated by nonlinear homeostatic variance.

**Figure 3.**
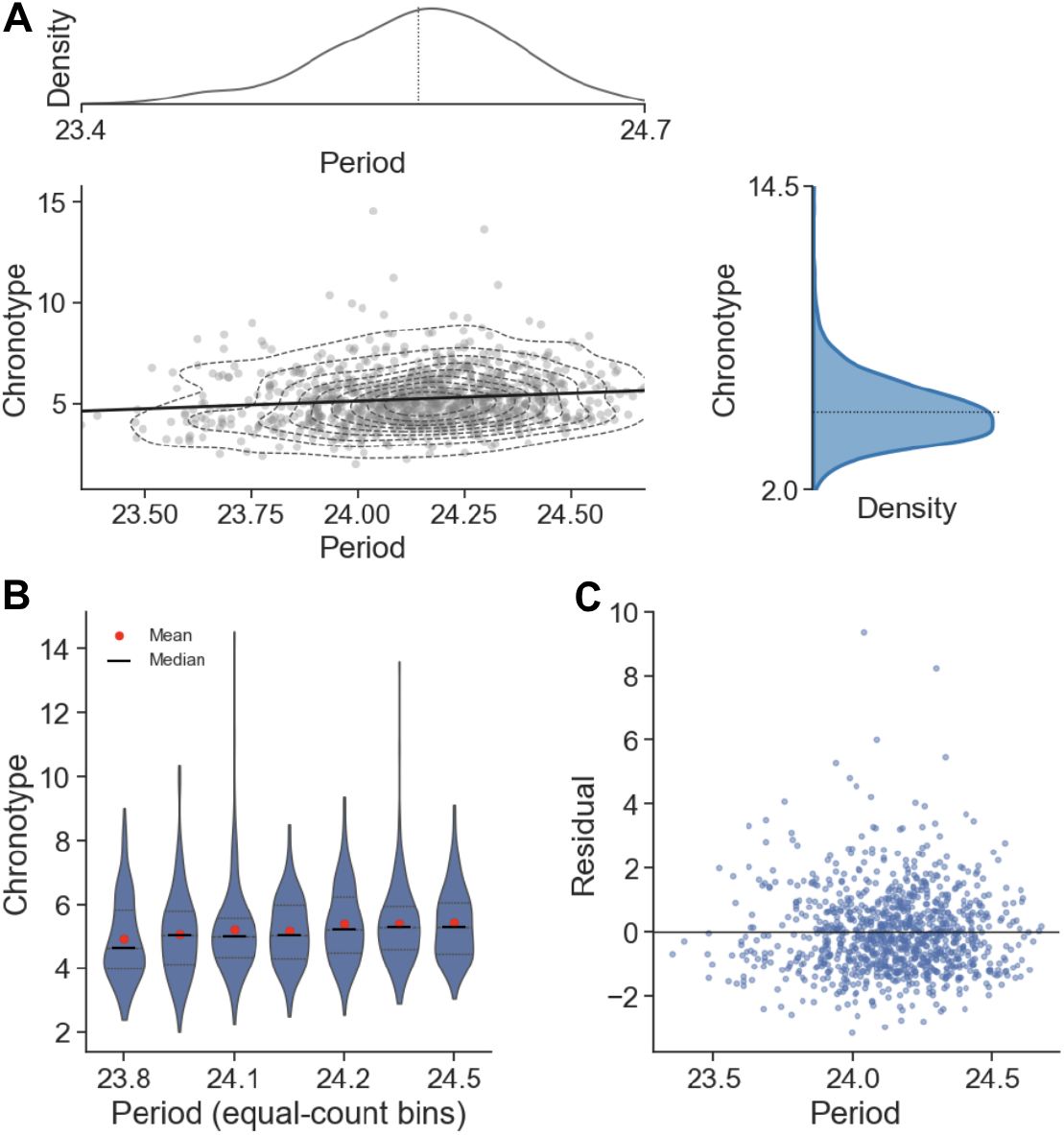
Relationship between intrinsic circadian period and chronotype. **(A)** Joint distribution of intrinsic circadian period (h) and chronotype (MSF, h) derived from the generative model. The solid line indicates the least-squares linear fit. The model reproduces the fundamental linearity observed in empirical data, and preserves the fundamental entrainment property where a longer period corresponds to a later chronotype. **(B)** Conditional distributions of chronotype visualized as violin plots across equal-count bins of circadian period. Red points indicate bin means and black bars indicate medians. **(C)** Residual analysis of the period-chronotype regression. The residuals exhibit a distinct asymmetric and nonconstant variance structure. The overall pattern mirrors the skewed heteroscedastic noise regime illustrated in Figure 1B, confirming that homeostatic sleep pressure introduces a structured deviation that transforms the left-skewed period input into a right-skewed behavioral output.

**Figure 4.**
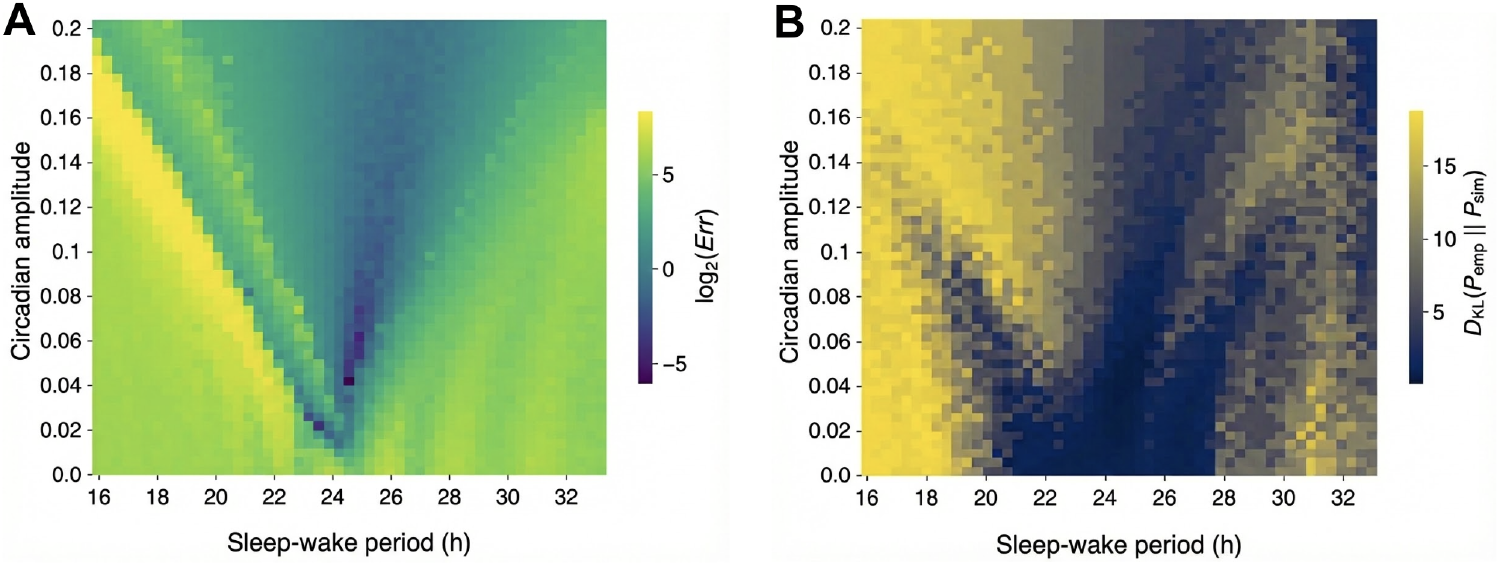
Dynamical stability and phenotypic plausibility across the parameter space. **(A)** Heatmap of the composite error function across sleep-wake period (*τ*, h) and circadian amplitude (*A*). Intrinsic sleep-wake period, the natural free-running frequency of the homeostatic oscillator in the absence of forcing. Circadian amplitude, the coupling strength of the circadian pacemaker (Process C). The error magnitude is plotted on a base-2 logarithmic scale (log_2_ *Err*). The low-error region forms a characteristic V-shaped Arnold tongue, delineating the boundaries of stable entrainment. High error regions (yellow) correspond to desynchronized states where phase drift results in a uniform distribution of sleep onsets, maximally deviating from empirical targets. **(B)** Heatmap of the KL divergence between simulated and empirical chronotype distributions. Unlike the stability landscape, the region of biological plausibility (low divergence) implies a non-monotonic relationship with forcing strength; it reaches maximal breadth at intermediate coupling (*A* = 0.1) before contracting at higher amplitudes (*A* = 0.2), indicating that excessive forcing suppresses the phase variance required to match human chronotype diversity.

### 3.3. Arnold tongue structure constrains the entrainment range for realistic chronotype distributions

To define the boundaries of stable sleep-wake regulation, we mapped the error landscapes for both the composite error function and the KL divergence across the parameter space of intrinsic sleep period and circadian amplitude (**Figure 4**). The stability landscape (**Figure 4A**) revealed the fundamental limits of entrainment, exhibiting a distinct Arnold tongue structure characteristic of coupled oscillators. This error function serves as a proxy condition for synchronization because of the statistical properties of circadian desynchrony. In a desynchronized (drifting) state, sleep onset times are distributed effectively uniformly across the 24-h cycle over the observation period. Such a distribution inherently tends toward a median of ∼12:00 and a skewness of zero. These values deviate greatly from the empirical targets of a 4:36 AM median and 0.6 skewness, resulting in a higher error score (equation 10). The minimization of this function therefore simultaneously satisfies dynamical stability, ensuring the sleep-wake cycle is entrained to its own circadian period. The region of low error therefore depicts the entrainment region. In the absence of circadian forcing (*A* = 0), the region of synchronization is extremely narrow (24.0-25.0 h). As the coupling strength increases, this region expands systematically into a V-shape, reaching maximal breadth at an amplitude of *A* = 0.2. The sharp transition from low to high error at the boundaries (yellow lines) delineates the critical bifurcation contour between stable frequency locking and phase drifting, confirming that the sleep-wake oscillator is entrained by the circadian rhythm. Matching only the mean and skewness is insufficient to characterize distributional realism, as visually similar skewness values can arise from fundamentally different distributional shapes. The KL divergence landscape (**Figure 4B**) therefore evaluates how closely simulated chronotype distributions match empirical data by quantifying similarity across the full distribution, rather than relying on summary statistics alone. Unlike the monotonic expansion seen in the stability landscape, the region of realistic chronotype distribution is nonlinear. The viable window expands initially with forcing widening from 20.0-28.0 h at *A* = 0 to a maximum range of 18.0-33.0 h at *A* = 0.1. However, further increasing the amplitude to *A* = 0.2 paradoxically constrained the viable range to 22.0-29.0 h. This contraction suggests that while strong circadian forcing guarantees stable entrainment (as shown in **Figure 4A**), it suppresses the natural phase variance required to reproduce the skewed chronotype distributions observed in human populations. The skewed chronotype distribution therefore is not solely a linear mapping of the circadian oscillator but an emergent property of entrainment and sleep homeostasis influence. Together, these results suggest that physiological sleep-wake cycles operate within a period range of 18 to 29 h, constrained by an optimal circadian amplitude ceiling of approximately 0.1.

## 4. Discussion

### From circadian phase to behavioral chronotype

The concept of chronotype was fundamentally defined as the phase of entrainment, that is, the stable timing relationship an organism’s internal clock establishes with the 24 h day. Pioneering work by Aschoff (1966) on chaffinches provided one of the first experimental demonstrations of this concept. He demonstrated that a bird with a short endogenous circadian period (< 24 h) began its daily activity before the lights came on, establishing a positive phase angle, while a bird with a long period (> 24 h) began its activity after the lights came on, establishing a negative phase angle. To stay synchronized with the light cue, the clock responds to the phase gap relative to the light-dark cycle by a daily phase shift, predicted by its phase response curve (PRC), that exactly corrects the mismatch between its intrinsic period and 24 h (Daan & Pittendrigh, 1976). Chronotype thus became understood as the behavioral manifestation of this stable phase relationship, measured with gold-standard markers for entrained circadian phase such as the DLMO.

On the other hand, real-world large-scale data assessed by behavioral questionnaires aim to create a score that correlates with the underlying circadian phase (Horne & Östberg, 1976), which was later refined into time-based metrics like the mid-point of sleep (Zavada et al., 2005). This practical convention introduces a theoretical gap in understanding precisely how the physiological phase of entrainment manifests as sleep-wake behavior. Our reappraisal of this gap contributes to understanding the nature of chronotype not as a pure readout of the central clock’s period, but as a multidimensional phenotype modulated through interactions with various internal and external time cues. We propose that this gap can be bridged by a hierarchical entrainment scheme.

### Hierarchical entrainment between the circadian clock and sleep

To understand this multidimensional phenotype, it is necessary to separate the clock from the sleep-wake cycle. Classical evidence from Aschoff (1965) illustrated complete desynchronization in a human subject, in which multiple internal rhythms became fully out of phase, and the circadian clocks outran the sleep-wake cycle. Czeisler et al. (1980) demonstrated that sleep duration can vary two-to three-fold depending on the circadian phase at which sleep is initiated, while the periodicity of hormonal secretion remains close within 24 h. Similarly, Hashimoto et al. (1997) measured longitudinal sleep and plasma melatonin rhythms in a sighted young man (21 years old) under isolated conditions, showing that while sleep-wake episodes follow the imposed schedule, melatonin levels continue to free-run with a period of 24.18 h. Furthermore, Yamanaka and Waterhouse (2016) showed that after an isolation protocol, the circadian rhythm of core body temperature maintains a period of approximately 25 h, whereas the sleep-wake cycle can lengthen beyond 30 h or shorten below 20 h.

These classical studies prove that the sleep-wake cycle and the circadian pacemaker are distinct oscillators. We therefore propose a hierarchical entrainment model in which sleep-wake behavior entrains to the circadian oscillator, which itself entrains to the light-dark cycle, extending beyond standard coupled oscillator theory (Kronauer, 1987; Strogatz, 1987; Strogatz & Stewart, 1993), where synchronization occurs in two distinct stages: first, the sleep-wake cycle entrains to the internal circadian oscillator (the SCN), and second, the SCN entrains to the 24-h light-dark cycle. There exists a supervenience relationship between the circadian clock and the external light-dark cycle, such that the daily alternation of light and darkness entrains the internal circadian oscillator in a unidirectional manner. The presence of an Arnold tongue in the light-driven circadian entrainment diagram demonstrates this dependency: the circadian clock’s phase and period become locked to the external cycle within a range determined by the strength of the photic input. Analogously, the observation that sleep phase forms its own Arnold tongue-like fitting structure with respect to the circadian phase indicates another level of supervenience, namely, that sleep timing supervenes on the circadian rhythm. Here, the circadian amplitude functions as the effective strength of the Zeitgeber that entrains the sleep-wake oscillator and establishes a hierarchical chain of dynamical dependencies extending from the daily light-dark cycle to the sleep rhythm.

### Homeostatic dynamics govern chronotype distribution shape

Within this hierarchical framework, our results demonstrate that the influence of the circadian period on behavioral chronotype is secondary to that of sleep homeostasis. Physiologically, the slow-wave activity (SWA) of non-REM sleep is generally considered one of the best biomarkers of homeostatic sleep pressure dissipation (Porkka-Heiskanen, 2013), and is closely related to physiological processes during sleep, such as growth hormone secretion and parasympathetic regulation (Dijk, 2009). From the approximately exponential decline of EEG power density in the slow-wave range during sleep, the decay time constant of sleep pressure can be estimated (Daan et al., 1984). By contrast, there is no consensus biomarker for sleep pressure during wakefulness, and its time constant is often approximated by fitting an exponential rise toward an asymptotic upper limit. Our model shows that the population chronotype distribution is primarily shaped not by the distribution of the circadian period, but by the distribution of this homeostatic sleep decay time constant. This finding suggests that the most critical step in determining behavioral chronotype is the direct interaction between the sleep homeostatic system and the central circadian clock.

This framework directly explains the nonlinear relationship observed between *τ* and behavioral chronotype. For individuals with a normal *τ* ∼ 24 h, the sleep-wake cycle is well within the wide, stable center of the Arnold tongue-like regime. In this robustly entrained state, the underlying positive correlation between a longer *τ* and a later chronotype is allowed to emerge, a finding consistent with both classical theory and modern human studies (Aschoff, 1966; Duffy et al., 2001). However, for individuals with very short or long periods, the sleep-oscillator system operates at the narrow, unstable edges of the regime. This weak entrainment is highly susceptible to noise and perturbations. Our results, which show both high variance in chronotype and a breakdown of the linear correlation between *τ* and chronotype in these groups, provide strong support for this model. This finding aligns with human experimental data demonstrating that individuals with intrinsic periods far from 24 h are unable to stably entrain to weak Zeitgeber cycles (Wright et al., 2001). It also provides a potential link to the asymmetric association with clinical pathology. Indeed, individuals with extreme delayed sleep timing are known to be at a vastly elevated risk for severe psychiatric comorbidities, including depression and bipolar disorder, and adverse physiological outcomes (Zou et al., 2022; Rahim et al., 2025; Zhang et al., 2022). In this unstable regime, the powerful influence of the sleep homeostatic system masks the weaker influence of the circadian period.

### Limitations

To map this interindividual variability to population-level distributions of chronotype, we utilized the mathematical two-process model. Research involving knockout in circadian clock genes provides further evidence that, while total sleep time may remain normal, the sleep-wake rhythm can be significantly altered. For example, *Rev-erbα* knockout mice exhibit fragmented sleep patterns, with sleep distributed throughout the day, despite maintaining a total sleep duration comparable to that of wild-type mice (Mang et al., 2016). Flip-flop switch models explain this sleep-wake behavior as mutual inhibition between wake-promoting monoaminergic nuclei of the ascending arousal system in the brainstem and hypothalamic nuclei and sleep-promoting neurons in the ventrolateral preoptic area (Saper et al., 2005). Building on this physiology, more detailed models include both circadian and homeostatic drives as inputs to the sleep switch (e.g., Phillips & Robinson, 2007; Postnova et al., 2012). However, the additional state variables and parameters required by such biological realism make them less tractable for mapping distributional characteristics at the population level.

Consequently, our first limitation is that the present study is built upon a gross simplification of a complex biological system, which posits that Process S and Process C are two independent processes. There is accumulating evidence for a bidirectional relationship between these systems (Deboer, 2018). For instance, high homeostatic sleep pressure from sleep deprivation has been shown to modulate the circadian system by attenuating the phase-shifting response to light (Burgess, 2010). Conversely, the circadian system gates the expression of sleep homeostasis, influencing the rate of build-up and decay of slow-wave activity (SWA) across the 24-h day (Daan et al., 1984). A related simplification concerns the threshold asymmetry used in our model. Flattening the lower threshold creates a shallower angle of intersection with the homeostatic decay curve. This increases the sensitivity of wake-onset timing to small inter-individual variations in homeostatic parameters, enabling the model to reproduce the broad, continuous distribution of chronotypes observed in human populations, rather than artificially clustering wake times. Biologically, this asymmetry reflects the distinct role of the circadian pacemaker in the “wake-maintenance zone” where it actively accumulates high homeostatic pressure in the evening to prevent premature sleep onset (Strogatz et al., 1987). Conversely, sleep termination is primarily driven by the decline in homeostatic sleep pressure, with a more subtle circadian gating, justifying a reduced amplitude for the lower threshold.

A second, and perhaps more central, limitation is our use of the mid-point of sleep as the definition of chronotype. While this is a standard and invaluable convention for large-scale studies (Roenneberg et al., 2003), it is a behavioral proxy, not the “ground truth” physiological phase of the central clock (i.e., the DLMO). As we have argued, this proxy is a composite signal, heavily influenced not only by the circadian system but also by the very homeostatic dynamics we are investigating, as well as by external factors like social schedules and individual light exposure patterns. This convention is the source of the theoretical gap that this study aims to address. What we reviewed, i.e., the nonlinear relationship between circadian period and chronotype, and the large influence of homeostasis, exists within the context of this behavioral definition. Therefore, our conclusions pertain specifically to the dynamics of the behavioral manifestation of chronotype. They provide a robust explanation for the timing of sleep as it is observed in real-world populations, but do not necessarily represent the pure, unmasked output of the SCN. These limitations highlight the necessity of our hierarchical model to bridge the gap between classical theory and complex, real-world data, and they pave the way for future models that could incorporate more complex, bidirectional interactions between the homeostatic and circadian systems.

## 5. Conclusion

Our stochastic extension of the two-process model resolves a fundamental paradox in sleep epidemiology. While the intrinsic circadian period is traditionally assumed to share a simple, linear relationship with chronotype, this assumption fails to explain the heavy right-skewed distributions of chronotype observed in global population data. Individuals with longer circadian periods fall on the theoretical boundaries of stable entrainment, i.e., the edges of the Arnold tongue. Daily fluctuations in sleep pressure disproportionately delay sleep onset, creating the heavy right tail seen in chronotype distributions. This work calls for a critical reexamination of how large-scale questionnaire data are interpreted. Behavioral chronotype must be recognized as a composite phenotype rather than a direct proxy for the central circadian clock, despite the apparent linear relationship. This distinction has direct clinical relevance, as chronotype-based classifications are increasingly linked to psychiatric and cardiometabolic risk profiles.

## Data availability statement

The simulation codes and parameter sweep results supporting the findings of this article are available at https://github.com/braintimelab/Stochastic-chronotype-model. Empirical data were digitized from published figures as cited in the text.

## Acknowledgements

This work was supported by the Higher Education Sprout Project from the Ministry of Education (MOE), Taiwan (DP2-TMU-113-N-06, DP2-TMU-114-N-06) and by the National Science and Technology Council (NSTC), Taiwan (113-2314-B-038-121, 114-2320-B-038-052-MY3).

## Notes

https://github.com/braintimelab/Stochastic-chronotype-model

### Competing Interest Statement

The authors have declared no competing interest.

